# Relationship between Anti-Spike Protein Antibody Titers and SARS-CoV-2 *In Vitro* Virus Neutralization in Convalescent Plasma

**DOI:** 10.1101/2020.06.08.138990

**Authors:** Eric Salazar, Suresh V. Kuchipudi, Paul A. Christensen, Todd N. Eagar, Xin Yi, Picheng Zhao, Zhicheng Jin, S. Wesley Long, Randall J. Olsen, Jian Chen, Brian Castillo, Christopher Leveque, Dalton M. Towers, Jason Lavinder, Jimmy D. Gollihar, Jose Cardona, Gregory C. Ippolito, Ruth H. Nissly, Ian M. Bird, Denver Greenawalt, Randall M. Rossi, Abinhay Gontu, Sreenidhi Srinivasan, Indira B. Poojary, Isabella M. Cattadori, Peter J. Hudson, Nicole Joselyn, Laura Prugar, Kathleen Huie, Andrew Herbert, David W. Bernard, John Dye, Vivek Kapur, James M. Musser

## Abstract

Newly emerged pathogens such as SARS-CoV-2 highlight the urgent need for assays that detect levels of neutralizing antibodies that may be protective. We studied the relationship between anti-spike ectodomain (ECD) and anti-receptor binding domain (RBD) IgG titers, and SARS-CoV-2 virus neutralization (VN) titers generated by two different *in vitro* assays using convalescent plasma samples obtained from 68 COVID-19 patients, including 13 who donated plasma multiple times. Only 23% (16/68) of donors had been hospitalized. We also studied 16 samples from subjects found to have anti-spike protein IgG during surveillance screening of asymptomatic individuals. We report a strong positive correlation between both plasma anti-RBD and anti-ECD IgG titers, and *in vitro* VN titer. Anti-RBD plasma IgG correlated slightly better than anti-ECD IgG titer with VN titer. The probability of a VN titer ≥160 was 80% or greater with anti-RBD or anti-ECD titers of ≥1:1350. Thirty-seven percent (25/68) of convalescent plasma donors lacked VN titers ≥160, the FDA-recommended level for convalescent plasma used for COVID-19 treatment. Dyspnea, hospitalization, and disease severity were significantly associated with higher VN titer. Frequent donation of convalescent plasma did not significantly decrease either VN or IgG titers. Analysis of 2,814 asymptomatic adults found 27 individuals with anti-RBD or anti-ECD IgG titers of ≥1:1350, and evidence of VN ≥1:160. Taken together, we conclude that anti-RBD or anti-ECD IgG titers can serve as a surrogate for VN titers to identify suitable plasma donors. Plasma anti-RBD or anti-ECD titer of ≥1:1350 may provide critical information about protection against COVID-19 disease.

## Introduction

The recently emerged SARS-CoV-2 novel coronavirus causing COVID-19 disease has spread globally and is now responsible for massive human morbidity and mortality. The pathogen was first documented to cause severe respiratory infections in humans in Wuhan, China, beginning in late December, 2019 (1). Soon thereafter, the SARS-CoV-2 virus was characterized as a member of the betacoronavirus genus and recognized to be related to several bat coronaviruses and Severe Acute Respiratory Syndrome (SARS-CoV) and Middle East Respiratory Syndrome (MERS) coronaviruses. SARS-CoV-2 spread was unusually rapid, and COVID-19 disease has now been reported in virtually all major population centers globally. In the United States, more than 1,500,000 COVID-19 cases have been documented and the virus has caused greater than 100,000 deaths nationwide. Many metropolitan regions have been especially affected, including but not limited to Seattle, New York City, Chicago, Miami, and Detroit (2).

Management of COVID-19 infection has predominantly involved aggressive support care. Various treatment approaches are being studied, including direct viral replication inhibition (3), anti-inflammatory drugs, and passive antibody therapies. Currently, the only available passive antibody therapy for COVID-19 patients is transfusion of convalescent plasma obtained from recovered patients. The therapy is safe, and clinical trials assessing treatment or prophylactic efficacy are underway (4). Clinical trials assessing the use of hyperimmune IgG may begin soon.

The Food and Drug Administration (FDA) has recommended (5) that convalescent plasma with a virus neutralizing antibody titer of ≥1:160 be used for therapeutic transfusion. However, assays to determine viral neutralizing antibody titers are not widely available, in part because they are labor intensive, cumbersome, and require a biosafety level 3 laboratory if live virus is used. Inasmuch as the VN titers in most donor plasma are not known prior to transfusion, a more facile method to identify suitable convalescent plasma donors is needed.

This is an especially pressing matter, since an increasing number of COVID-19 patients are being treated globally with convalescent plasma. For example, under an FDA-approved expanded access protocol, nearly 20,000 transfusions have already occurred in the United States (6).

The trimeric spike (S) protein made by SARS-CoV-2 is a large molecule that is critical to virus dissemination and pathogenesis. S protein is a densely glycosylated molecule present on the surface of the virus. S protein mediates binding of the SARS-CoV-2 virus to the host angiotensin-converting enzyme 2 (ACE2), thereby acting as the first step in cell entry and infection. Recent work has shown that SARS-CoV-2 and SARS-CoV-1 share the same ACE2 receptor. The molecular mechanism used by S protein to gain entry into host cells is complex and involves a region of the molecule known as the receptor binding domain (RBD). Engagement of S protein with the host receptor results in considerable changes in molecular conformation. The S protein has a critical function in host-cell entry, and thus is a major target for vaccine research and antibody-mediated VN efforts.

Many lines of evidence from studies of SARS-CoV-1, MERS, and SARS-CoV-2 show that infected hosts make antibody directed against S protein (7–14). In addition, immunization with S protein can protect laboratory animals against experimental infection with SARS-CoV-1, MERS-CoV, and SARS-CoV-2 (15–19). Similarly, IgG directed against S protein has been reported to have *in vitro* VN activity.

The goal of this study was to test the hypothesis that anti-ECD and/or anti-RBD IgG titer are correlated with VN titer, and thus could be used as a surrogate marker to identify plasma donors with titers above the FDA threshold value of 1:160. To test this hypothesis, we studied plasma and serum samples from 68 recovered COVID-19 patients with documented disease based on a positive molecular test for SARS-CoV-2. VN titer was determined independently in two laboratories using two different *in vitro* assays. The results show a strong positive correlation between anti-RBD and anti-ECD plasma IgG ELISA titers and the magnitude of *in vitro* VN. Specifically, we report that there is an 80% probability or greater of a VN titer at or above the FDA recommended level of 1:160 for COVID-19 convalescent plasma with anti-RBD or anti-ECD IgG titers of ≥1:1350. The results provide an important quantitative target for therapeutic and prophylactic treatments. We also find that convalescent donors maintain high-titer anti-RBD and anti-ECD IgG with *in vitro* VN activity over many weeks. Frequent plasma donations do not cause a significant decrease in antibody or VN titers. Finally, analysis of anti-ECD and anti-RBD IgG titers in 2,814 asymptomatic individuals in a surveillance cohort identified 27 individuals with anti-RBD or anti-ECD IgG titers of ≥1:1350, and evidence of VN ≥1:160. Thus, some asymptomatic individuals may have plasma suitable for therapeutic use and may have a degree of relative immunity against SARS-CoV-2.

## Results

### Plasma Donor Characteristics

Ninety-three samples from 68 unique COVID-19 convalescent plasma donors were assessed (Table 1). The average age was 45 (range 23 to 78) and 36 donors were female. Most donors had mild to moderate disease, with 44% (30/68) having a symptom severity score of 1,32% (22/68) having score of 2, 10% (7/68) having a score of 3, 7% (5/68) having a score of 4, and 6% (4/68) having a score of 5. Sixteen donors required hospitalization, with an average length of stay of 4 d (range 2-13 d). Thirteen individuals donated more than once (range 1-7 times) with most (9/13) donating twice only. For all samples assessed, the median interval from symptom onset to donation visit was 32 d (range 17-53 d; IQR 28-36 d), and the median interval from symptom resolution to donation visit was 20 d (range 15-38 d; IQR 17-25 d).

**Table 1:** Demographics and Characteristics of Convalescent Plasma Donors and Samples.

| Sample # | Subject ID | Gender | Age | Severity | Hospitalized? | LOS (Days) | Symptom Duration (Days) | Days Since Symptom Onset | Days Since Symptom Resolution | Visit # | ECD (Titer) | RBD (Titer) | VN (Titer) | VN2 (IC50) | Clade |
| --- | --- | --- | --- | --- | --- | --- | --- | --- | --- | --- | --- | --- | --- | --- | --- |
| 1 | HMH0001 | M | 44 | 1 | NO | NA | 3 | 20 | 17 | 1 | 150 | 1350 | 320 | 53.2 | A2a |
| 2 | HMH0001 | M | 44 | 1 | NO | NA | 3 | 24 | 21 | 2 | NA | NA | 640 | 48.6 | A2a |
| 3 | HMH0001 | M | 44 | 1 | NO | NA | 3 | 27 | 24 | 3 | 150 | 450 | 320 | 53.3 | A2a |
| 4 | HMH0001 | M | 44 | 1 | NO | NA | 3 | 31 | 28 | 4 | 150 | 800 | 320 | 46.8 | A2a |
| 5 | HMH0001 | M | 44 | 1 | NO | NA | 3 | 34 | 31 | 5 | 150 | 800 | 320 | 96.3 | A2a |
| 6 | HMH0001 | M | 44 | 1 | NO | NA | 3 | 38 | 35 | 6 | 450 | 450 | 160 | 69.4 | A2a |
| 7 | HMH0001 | M | 44 | 1 | NO | NA | 3 | 41 | 38 | 7 | 450 | 450 | 80 | 71.5 | A2a |
| 8 | HMH0002 | M | 54 | 1 | NO | NA | 13 | 28 | 15 | 1 | 50 | 150 | 40 | 1.5 | A2a |
| 9 | HMH0003 | M | 36 | 1 | NO | NA | 7 | 25 | 18 | 1 | 450 | 1350 | 80 | 38.4 | B |
| 10 | HMH0003 | M | 36 | 1 | NO | NA | 7 | 28 | 21 | 2 | 150 | 450 | 80 | 10.4 | B |
| 11 | HMH0003 | M | 36 | 1 | NO | NA | 7 | 33 | 26 | 3 | 450 | 800 | 80 | 7.6 | B |
| 12 | HMH0003 | M | 36 | 1 | NO | NA | 7 | 35 | 28 | 4 | 450 | 450 | 0 | 8 | B |
| 13 | HMH0003 | M | 36 | 1 | NO | NA | 7 | 40 | 33 | 5 | 450 | 450 | 320 | 19.8 | B |
| 14 | HMH0003 | M | 36 | 1 | NO | NA | 7 | 42 | 35 | 6 | 150 | 450 | 20 | 1.6 | B |
| 15 | HMH0004 | F | 54 | 2 | NO | NA | 18 | 32 | 14 | 1 | 1350 | 1350 | 320 | 270.7 | NA |
| 16 | HMH0004 | F | 54 | 2 | NO | NA | 18 | 36 | 18 | 2 | 1350 | 1350 | 640 | 128.5 | NA |
| 17 | HMH0009 | F | 38 | 2 | NO | NA | 12 | 30 | 18 | 1 | 450 | 450 | 80 | 4.8 | NA |
| 18 | HMH0011 | F | 67 | 1 | NO | NA | 11 | 28 | 17 | 1 | 0 | 50 | 40 | 8 | A2a |
| 19 | HMH0012 | F | 46 | 1 | NO | NA | 16 | 30 | 14 | 1 | 150 | 450 | 320 | 30 | NA |
| 20 | HMH0013 | F | 43 | 1 | NO | NA | 11 | 28 | 17 | 1 | 1350 | 3200 | 320 | 214.2 | A2a |
| 21 | HMH0016 | F | 47 | 1 | NO | NA | 13 | 32 | 19 | 1 | 4050 | 4050 | 320 | 234.2 | A2a |
| 22 | HMH0020 | F | 41 | 2 | NO | NA | 2 | 17 | 15 | 1 | 50 | 200 | 20 | NA | NA |
| 23 | HMH0028 | M | 23 | 1 | NO | NA | 12 | 31 | 19 | 1 | 150 | 150 | 20 | 0.6 | A2a |
| 24 | HMH0029 | F | 66 | 1 | NO | NA | 6 | 22 | 16 | 1 | 150 | 450 | 80 | 9.3 | A2a |
| 25 | HMH0032 | M | 65 | 2 | NO | NA | 11 | 25 | 14 | 1 | 450 | 4050 | 320 | 94.8 | NA |
| 26 | HMH0035 | M | 50 | 2 | NO | NA | 14 | 28 | 14 | 1 | 1350 | 3200 | 320 | 47.3 | B |
| 27 | HMH0035 | M | 50 | 2 | NO | NA | 14 | 38 | 24 | 2 | 1350 | 1350 | 640 | 72 | B |
| 28 | HMH0040 | M | 52 | 2 | NO | NA | 12 | 29 | 17 | 1 | NA | NA | 1280 | 417 | A2a |
| 29 | HMH0040 | M | 52 | 2 | NO | NA | 12 | 35 | 23 | 2 | 4050 | 4050 | 640 | 724.5 | A2a |
| 30 | HMH0040 | M | 52 | 2 | NO | NA | 12 | 37 | 25 | 3 | 1350 | 4050 | 320 | 274 | A2a |
| 31 | HMH0045 | F | 23 | 1 | NO | NA | 9 | 33 | 24 | 1 | 4050 | 1350 | 1280 | 63.8 | A2a |
| 32 | HMH0049 | F | 57 | 1 | NO | NA | 12 | 27 | 15 | 1 | 150 | 450 | 320 | 47 | NA |
| 33 | HMH0050 | M | 41 | 2 | NO | NA | 7 | 30 | 23 | 1 | 1350 | 1350 | 320 | 18.9 | NA |
| 34 | HMH0050 | M | 41 | 2 | NO | NA | 7 | 33 | 26 | 2 | 150 | 150 | 320 | 21.4 | NA |
| 35 | HMH0051 | F | 50 | 1 | NO | NA | 16 | 30 | 14 | 1 | 150 | 450 | 160 | 58.4 | NA |
| 36 | HMH0052 | F | 27 | 3 | YES | 2 | 17 | 31 | 14 | 1 | 1350 | 1350 | 160 | 100.1 | A2a |
| 37 | HMH0053 | M | 29 | 2 | NO | NA | 10 | 28 | 18 | 1 | 450 | 450 | 160 | 18.9 | NA |
| 38 | HMH0055 | M | 61 | 3 | YES | 3 | 14 | 33 | 19 | 1 | 1350 | 3200 | 320 | 134.1 | B |
| 39 | HMH0057 | F | 44 | 2 | NO | NA | 6 | 34 | 28 | 1 | 450 | 450 | 160 | 111.5 | A2a |
| 40 | HMH0062 | F | 24 | 1 | NO | NA | 13 | 32 | 19 | 1 | 1350 | 1350 | 10 | 443.9 | A2a |
| 41 | HMH0062 | F | 24 | 1 | NO | NA | 13 | 35 | 22 | 2 | 1350 | 1350 | 1280 | 288.8 | A2a |
| 42 | HMH0069 | F | 49 | 1 | NO | NA | 6 | 28 | 22 | 1 | 450 | 450 | 80 | NA | A2a |
| 43 | HMH0069 | F | 49 | 1 | NO | NA | 6 | 32 | 26 | 2 | 450 | 450 | 80 | 35.3 | A2a |
| 44 | HMH0070 | F | 37 | 2 | NO | NA | 19 | 38 | 19 | 1 | 450 | 1350 | 160 | 126.7 | A2a |
| 45 | HMH0072 | F | 23 | 1 | NO | NA | 13 | 29 | 16 | 1 | 0 | 0 | 0 | NA | NA |
| 46 | HMH0072 | F | 23 | 1 | NO | NA | 13 | 37 | 24 | 2 | 50 | 50 | 0 | NA | NA |
| 47 | HMH0088 | F | 29 | 1 | NO | NA | 3 | 21 | 18 | 1 | 0 | 50 | 20 | NA | A2a |
| 48 | HMH0089 | F | 42 | 2 | NO | NA | 18 | 38 | 20 | 1 | 450 | 450 | 160 | 95 | A2a |
| 49 | HMH0090 | M | 33 | 1 | NO | NA | 3 | 37 | 34 | 1 | 150 | 150 | 1280 | 51.3 | A2a |
| 50 | HMH0099 | F | 54 | 1 | NO | NA | 1 | 20 | 19 | 1 | 50 | 50 | 10 | NA | NA |
| 51 | HMH0112 | F | 47 | 1 | NO | NA | 1 | 32 | 31 | 1 | 450 | 450 | 40 | 78.9 | A2a |
| 52 | HMH0113 | F | 52 | 1 | NO | NA | 9 | 29 | 20 | 1 | 1350 | 150 | 40 | 6.9 | NA |
| 53 | HMH0116 | M | 27 | 1 | NO | NA | 16 | 32 | 16 | 1 | 450 | 450 | 20 | 11.3 | NA |
| 54 | HMH0117 | F | 27 | 2 | NO | NA | 10 | 30 | 20 | 1 | 1350 | 1350 | 320 | 32.7 | NA |
| 55 | HMH0118 | F | 50 | 1 | NO | NA | 15 | 34 | 19 | 1 | 1350 | 450 | 320 | 8 | NA |
| 56 | HMH0119 | F | 35 | 1 | NO | NA | 6 | 25 | 19 | 1 | 450 | 450 | 0 | 47.3 | NA |
| 57 | HMH0120 | F | 41 | 1 | NO | NA | 3 | 19 | 16 | 1 | 450 | 800 | 320 | 62.3 | NA |
| 58 | HMH0121 | F | 51 | 1 | NO | NA | 7 | 21 | 14 | 1 | 150 | 200 | 40 | 25.4 | NA |
| 59 | HMH0133 | M | 51 | 2 | NO | NA | 4 | 25 | 21 | 1 | 150 | 450 | 320 | 138.8 | NA |
| 60 | HMH0135 | F | 47 | 2 | NO | NA | 12 | 32 | 20 | 1 | 4050 | 4050 | 320 | 174.4 | NA |
| 61 | HMH0137 | F | 53 | 1 | NO | NA | 8 | 38 | 30 | 1 | 450 | 800 | 0 | 2.5 | NA |
| 62 | HMH0143 | F | 49 | 2 | NO | NA | 10 | 37 | 27 | 1 | 1350 | 3200 | 160 | 72.4 | NA |
| 63 | HMH0144 | M | 48 | 5 | YES | 3 | 15 | 40 | 25 | 1 | 1350 | 3200 | 640 | 723 | NA |
| 64 | HMH0144 | M | 48 | 5 | YES | 3 | 15 | 45 | 30 | 2 | 4050 | 1350 | 640 | 423.1 | NA |
| 65 | HMH0144 | M | 48 | 5 | YES | 3 | 15 | 50 | 35 | 3 | 450 | 1350 | 640 | 453.8 | NA |
| 66 | HMH0144 | M | 48 | 5 | YES | 3 | 15 | 53 | 38 | 4 | 4050 | 4050 | 1280 | 278.3 | NA |
| 67 | HMH0156 | M | 59 | 1 | NO | NA | 6 | 22 | 16 | 1 | 4050 | 1350 | 1280 | 814.8 | NA |
| 68 | HMH0156 | M | 59 | 1 | NO | NA | 6 | 29 | 23 | 2 | 1350 | 1350 | 1280 | 751.7 | NA |
| 69 | HMH0158 | M | 33 | 2 | NO | NA | 10 | 26 | 16 | 1 | 150 | 200 | 40 | 33.8 | NA |
| 70 | HMH0162 | F | 51 | 3 | YES | 3 | 9 | 34 | 25 | 1 | 150 | 450 | 160 | 28.2 | NA |
| 71 | HMH0229 | M | 32 | 2 | NO | NA | 21 | 40 | 19 | 1 | 450 | 1350 | 80 | 311.3 | NA |
| 72 | HMH0234 | M | 40 | 2 | NO | NA | 12 | 27 | 15 | 1 | 150 | 150 | 40 | 25 | NA |
| 73 | HMH0245 | M | 51 | 5 | YES | 4 | 14 | 38 | 24 | 1 | 1350 | 4050 | 320 | 145 | NA |
| 74 | HMH0249 | M | 56 | 1 | NO | NA | 8 | 22 | 14 | 1 | 4050 | 4050 | 320 | 415.8 | NA |
| 75 | HMH0255 | M | 40 | 2 | NO | NA | 14 | 31 | 17 | 1 | 450 | 450 | 40 | 22 | NA |
| 76 | HMH0260 | M | 44 | 2 | NO | NA | 5 | 24 | 19 | 1 | 4050 | 1350 | 1280 | 468.5 | NA |
| 77 | HMH0262 | F | 36 | 4 | YES | 2 | 16 | 31 | 15 | 1 | 4050 | 4050 | 1280 | 262.8 | A2a |
| 78 | HMH0265 | F | 53 | 2 | NO | NA | 11 | 31 | 20 | 1 | 4050 | 4050 | 320 | 188 | NA |
| 79 | HMH0313 | M | 78 | 3 | YES | 1 | 16 | 36 | 20 | 1 | 4050 | 4050 | 160 | 77.8 | NA |
| 80 | HMH0363 | M | 56 | 1 | NO | NA | 7 | 34 | 27 | 1 | 450 | 1350 | 80 | 26.4 | NA |
| 81 | HMH0368 | F | 37 | 2 | NO | NA | 9 | 29 | 20 | 1 | 450 | 1350 | 160 | 65.4 | NA |
| 82 | HMH0369 | M | 41 | 1 | NO | NA | 21 | 39 | 18 | 1 | 150 | 450 | 80 | 56.1 | NA |
| 83 | HMH0376 | M | 52 | 4 | YES | 7 | 14 | 28 | 14 | 1 | 1350 | 4050 | 1280 | 74.8 | NA |
| 84 | HMH0376 | M | 52 | 4 | YES | 7 | 14 | 32 | 18 | 2 | 4050 | 4050 | 160 | 90.7 | NA |
| 85 | HMH0430 | M | 44 | 4 | YES | 4 | 14 | 35 | 21 | 1 | 4050 | 4050 | 1280 | 207.4 | NA |
| 86 | HMH0576 | F | 50 | 3 | YES | 5 | 2 | 41 | 39 | 1 | 0 | 0 | 0 | NA | NA |
| 87 | HMH0580 | F | 43 | 3 | YES | 4 | 12 | 29 | 17 | 1 | 4050 | 4050 | 1280 | 752.3 | A2a |
| 88 | HMH0580 | F | 43 | 3 | YES | 4 | 12 | 35 | 23 | 2 | 1350 | 1350 | 1280 | 421.4 | A2a |
| 89 | HMH0598 | M | 46 | 4 | YES | 6 | 29 | 30 | 1 | 1 | 4050 | 4050 | 1280 | 365.7 | NA |
| 90 | HMH0620 | M | 59 | 5 | YES | 6 | 13 | 40 | 27 | 1 | 4050 | 4050 | 1280 | 645.9 | NA |
| 91 | HMH0634 | M | 53 | 5 | YES | 13 | 16 | 33 | 17 | 1 | 4050 | 4050 | 640 | 148.1 | B |
| 92 | HMH0699 | M | 61 | 3 | YES | 2 | 5 | 35 | 30 | 1 | 4050 | 450 | 640 | 343 | NA |
| 93 | HMH0879 | M | 50 | 4 | YES | 5 | 14 | 34 | 20 | 1 | 4050 | 4050 | 160 | 178.1 | A2a |

Abbreviations: F, Female; M, Male; LOS, Length of Stay; ECD, anti-spike ectodomain; RBD, Receptor Binding Domain; VN, Virus Neutralization

We also studied plasma from 16 asymptomatic individuals identified during an institutional surveillance program involving 2,814 individuals (manuscript in preparation). Of these 2,814 individuals, 102 had ELISA titers ≥50 (Supplemental Fig. 2). The average age of these 16 asymptomatic donors was 35 (range 20-49) and half were female (Supplemental Table 1).

### VN Titers in Convalescent Plasma Donors

VN titers in samples from COVID-19 convalescent plasma donors were assessed with a traditional microneutralization assay evaluating protection from virus infection as determined by crystal violet staining 3 d post-infection. Plasma samples from the majority of donors (43/68; 63%) had a VN titer ≥1:160, the FDA recommended VN antibody titer for convalescent plasma to be used for therapeutic transfusion purposes. In contrast, 25 of 68 donors (37%) had a plasma titer below this recommended cutoff value (Figure 1A and Table 1).

**Fig 1.**
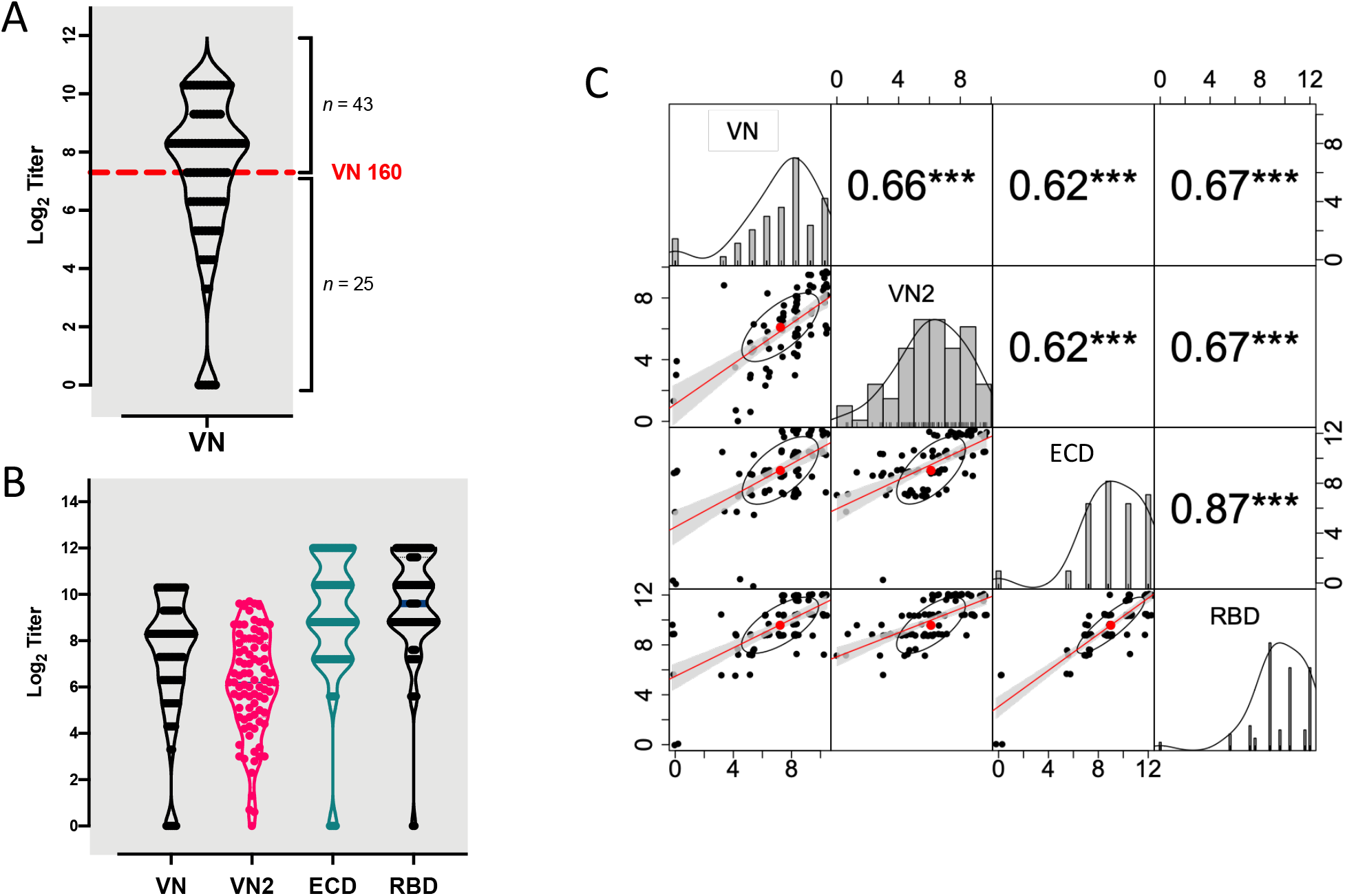
Patterns of VN and ELISA titers. A) Violin plot of distribution of VN titers at initial donation. Number of donor cases (total = 68) above and below the VN 160 cut off value are reported. B) Violin plots showing similar patterns of distribution of titers at initial donation for the two VN assays, together with the reciprocal ELISA IgG titers for plasma anti-ECD protein (ECD) and anti-RBD IgG (RBD). C) Pair-wise Pearson correlations showing the correlation coefficient (r) and related significant value (***= p<0.001) above the diagonal, and the bivariate scatterplots (jittered points) with linear regression fit (red line), confidence intervals (grey shading), correlation value (red point) and correlation ellipse (black ellipse) below the diagonal. The density plot (black line) and histogram of each variable is reported along the diagonal. Data are presented in log2-scale of reciprocal titers for VN, anti-ECD IgG and anti-RBD IgG, and in IC50 units for VN2.

### Correlation between Two VN Assays

VN titers were assessed blinded (that is, without knowledge of the data generated by laboratory one) in a second laboratory with a different microneutralization assay (VN2) that determined the percentage of infected cells 24 h post-infection using a SARS-CoV-S specific mAb and a fluorescently labeled secondary antibody. The results from the two VN assays were highly correlated (r=0.66, P < 0.001) (Figure 1B and C).

### Association between ELISA IgG Titers and VN Titer

Recognizing the urgent need for assays that could serve as a surrogate for VN, we assessed the association between ELISA anti-ECD and anti-RBD IgG titers and VN titers. The results of all four assays (anti-ECD and anti-RBD ELISAs, VN, and VN2) were strongly correlated (Figure 1C). Anti-RBD IgG had a numerically but not statistically greater correlation than anti-ECD (0.67 versus 0.62) with both microneutralization assays. We found that more than 80% of donors had a VN titer ≥1:160 in convalescent plasma when their serum anti-RBD or anti-EDC titers were 1:1350 or higher (Figure 2). Importantly, samples from naïve human plasma specimens obtained before the discovery of SARS-CoV-2 had no detectable titer in any of the four assays (data not shown).

**Fig 2.**
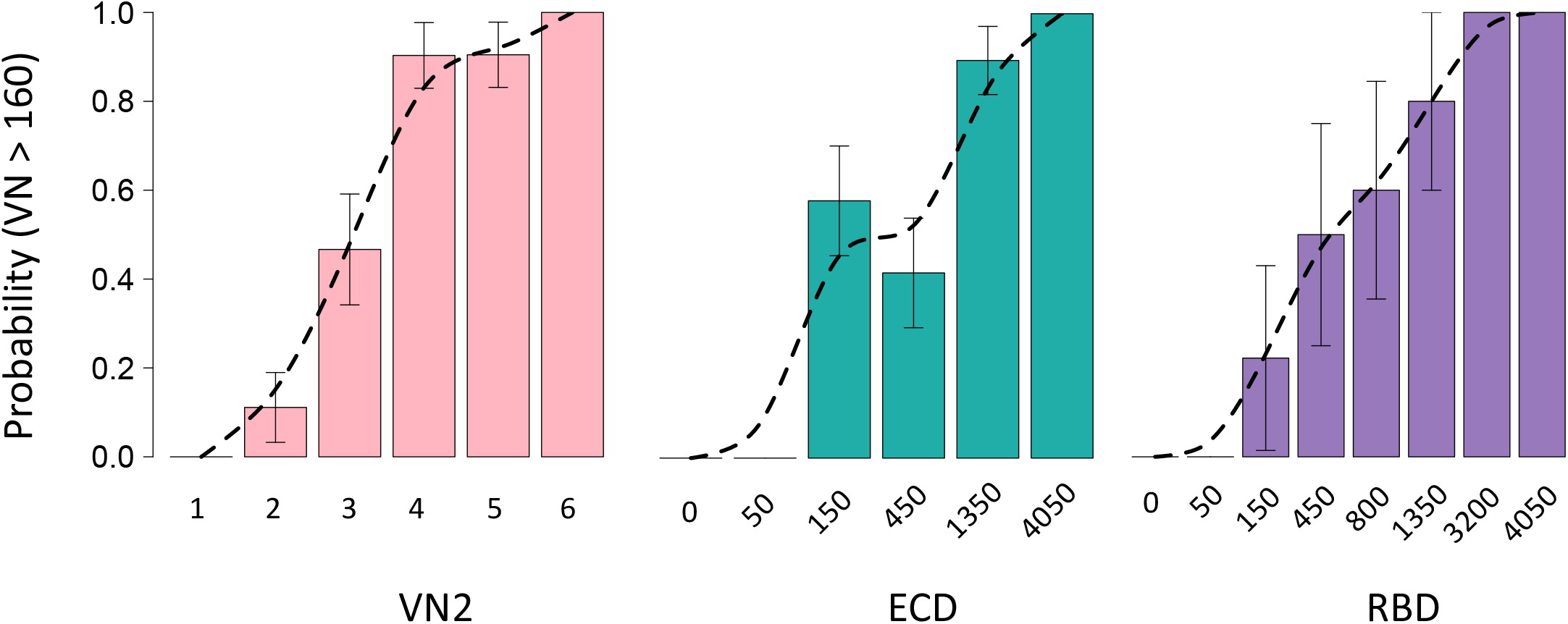
Bar plots reporting the prevalence of donors with VN>160 for VN2, ECD or RBD. Probabilities of VN160 were plotted for six range classes, with an interclass interval of 1.8 log2 IC50 values (class 1 - <2; class 2 - 2,12; class 3-12,42; class 4 - 42-147; class 5 - 147,512; class 6 - >512) or observed classes for ECD (*n* = 6) and RBD (*n* = 8) reciprocal ELISA titers. A spline curve (dotted line, smoothness shape=1) has been fitted to the probability values and standard errors (bars) are reported.

### Relationship between Antibody Titers and Donor Characteristics

Inasmuch as approximately one-third of donors lacked convalescent plasma with the FDA-recommended VN titer cutoff of ≥1:160, we sought to identify donor characteristics that may be associated with a higher IgG titer. Such characteristics could aid donor recruitment efforts by identifying which recovered patients may have mounted a strong humoral response. We found that the presence of dyspnea during COVID-19 disease, hospitalization requirement, and more severe disease are all positively and significantly associated with higher IgG titers in all assays (Figure 3). Duration of disease symptoms was not associated with titer. Nor was there an association with time of plasma collection, since symptom onset and titer in the donor population occurred more than 14 days after symptom resolution (as required by the FDA). These results suggest that donors had already plateaued in their IgG titer at the time plasma was obtained, as there was no appreciable trend in titer increase over time (Supplemental Figure 1A and B).

**Fig 3.**
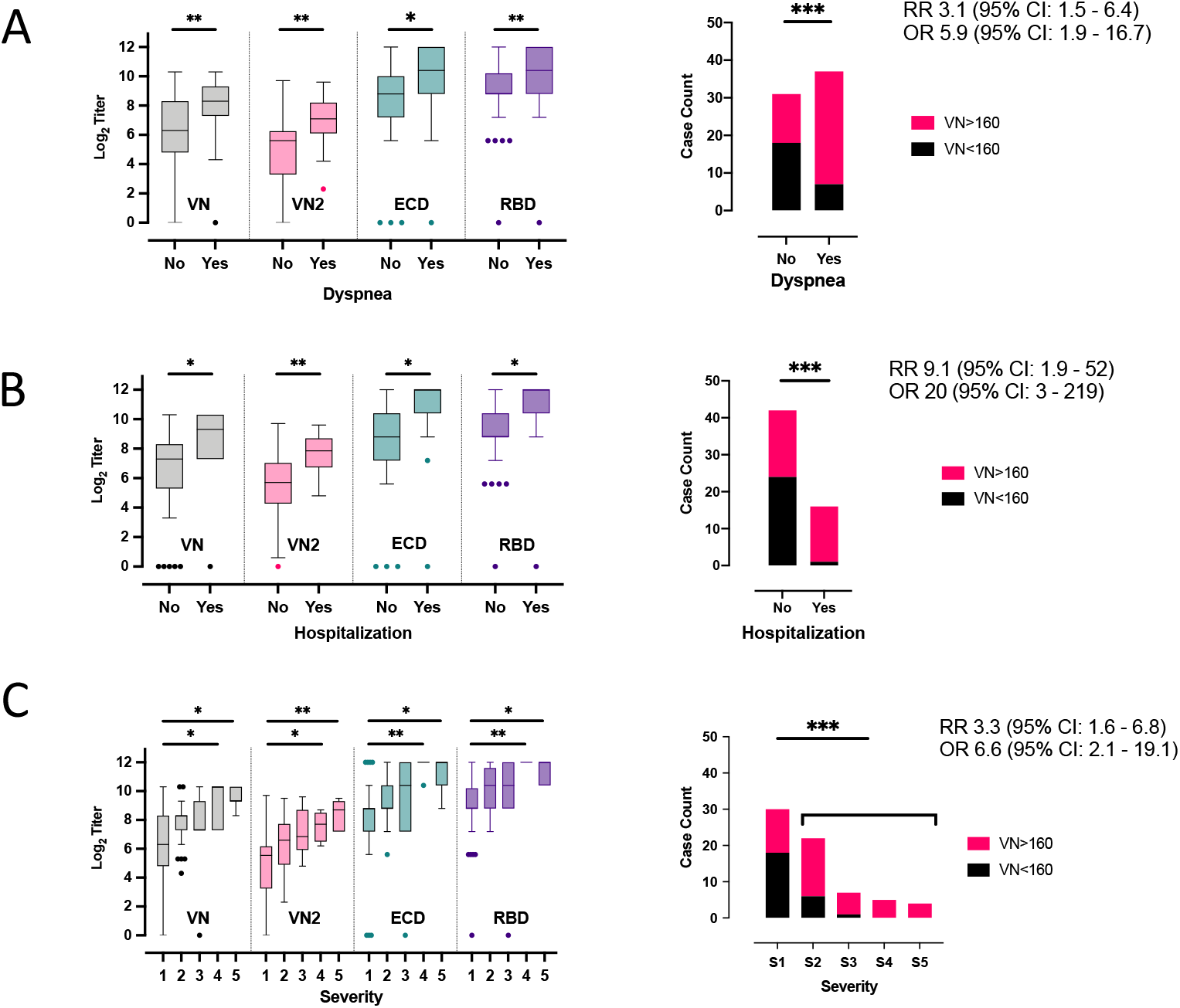
Boxplots of VN, VN2, anti-ECD and anti RBD titers by A) dyspnea, B) hospitalization and C) disease severity (1= low severity, 5 high severity) at initial plasma donation from the 68 individual donors. The median, minimum, maximum, first and third quartile and extreme values are reported. Barplots showing case counts of donors above and below the VN 160 threshold stratified by whether they self-reported A) occurrence of dyspnea during symptomatic phase of disease; B) hospitalization; and, C) disease severity. Pair-wise t-test (significant comparisons (*=p<0.05, **= p<0.01, ***= p<0.001), odds ratio (OR) and relative risk (RR) with confidence intervals (CI) are also reported.

**Fig 4.**
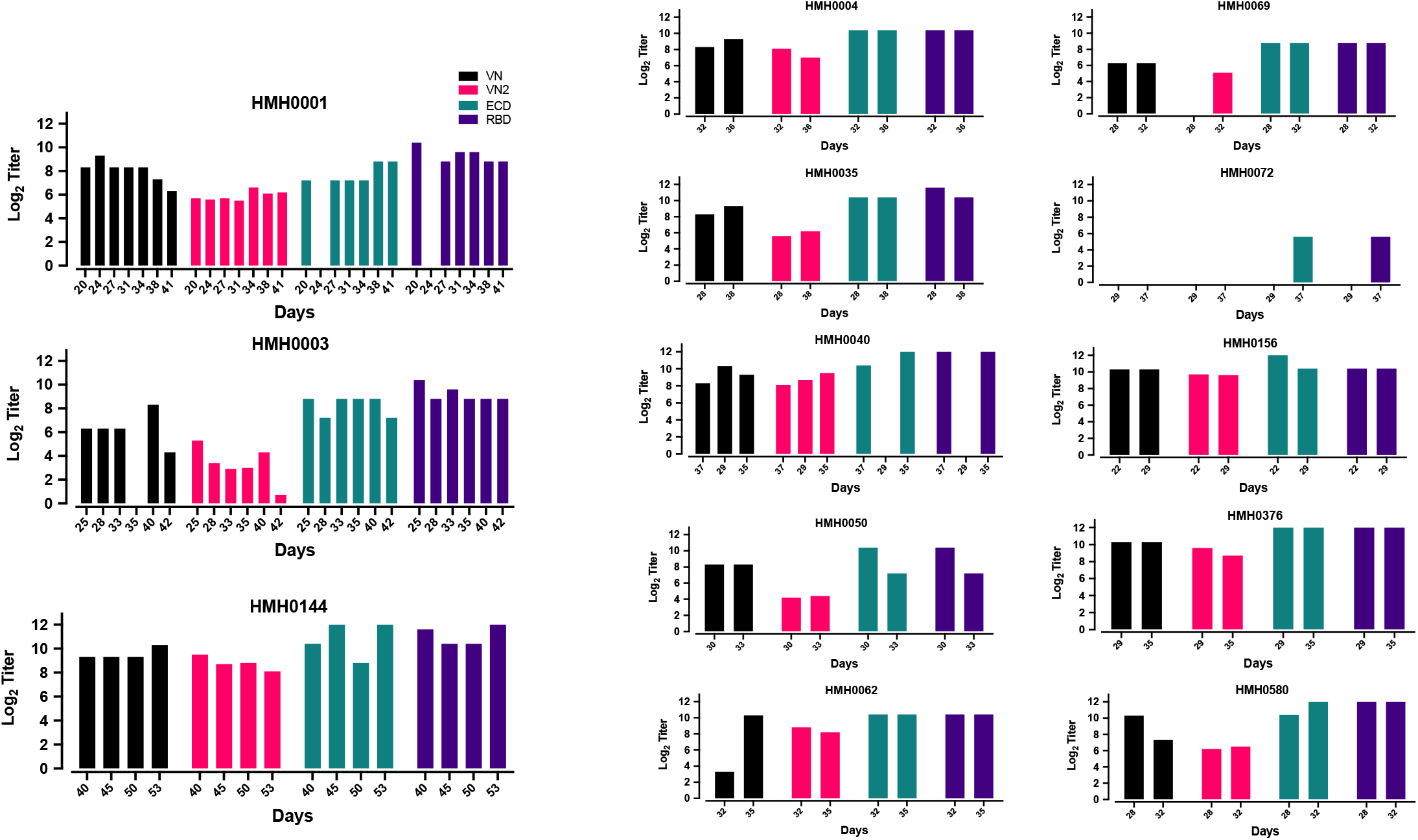
Trends in VN, VN2, anti-ECD IgG and anti-RBD IgG titers for donors with multiple consecutive donations. VN, ECD, and RBD are reported as log 2 of reciprocal titers whereas VN2 is represented by the log2 of IC50 value. HMH number refers to arbitrary number assigned to each plasma donor.

### VN Titers Over Time from the Same Convalescent Plasma Donors

Thirteen individuals donated convalescent plasma more than once (range, 2 to 7 donations). The availability of longitudinal samples from the same plasma donors permitted us to assess the arc of anti-ECD and anti-RBD IgG titers and VN over time within individuals. There was no significant decrease in IgG titers as assessed by the ELISA or VN titer (Supplemental Figure 2), even among donors who donated twice/week for up to seven donations. Thus, we observed stable high titers both within and between individual donors.

### Relationship between Infecting Strain Clade and VN Titer

We had available the virus genome sequences obtained from clinical samples (e.g., nasopharyngeal swab, oropharyngeal swab, or sputum) from 25 plasma donors. Eighty-four percent (21/25) of donors had been infected with strain A2a, and the remaining donors had been infected with strain B. Although the number of specimens is small, we tested the hypothesis that a relationship exists between the VN titer and genetic clade of the infecting SARS-CoV-2 strain. No definitive relationship was evident from analysis of the available data (Table 1).

### Asymptomatic Individuals and VN Titers

Having established a relationship between IgG titer and *in vitro* SARS-CoV-2 VN titer, we next determined IgG titers in a sample of 2,814 asymptomatic adults screened under a surveillance protocol. We found that 67 of 2,814 (2.4%) individuals had anti-ECD and anti-RBD IgG ELISA titers of ≥1:150, of which 27 had anti-RBD or anti-ECD IgG titers of ≥1350. Among 16 of the 67 specimens chosen for VN analysis based on a range of titers [150 (*n* = 5), 450 (*n* = 5), and 1350 (*n* = 6)], seven had a VN titer ≥160. All individuals with a VN titer ≥160 had anti-RBD IgG titers of ≥1350. Conversely, with a single exception, all individuals with an anti-RBD IgG titer of ≥1350 had a VN titer ≥160 (Supplemental Table 1). Overall, among the 16 samples tested in this cohort, anti-RBD and anti-ECD ELISA titers also were strongly correlated with *in vitro* VN titer (R > 0.75, P = 0.001).

## Discussion

In the absence of an efficacious vaccine to prevent COVID-19 disease, there is a pressing need for assays that detect neutralizing antibodies against SARS-CoV-2. Here we studied the relationship between anti-RBD and anti-ECD IgG titers present in convalescent plasma obtained from COVID-19 patients and *in vitro* SARS-CoV-2 VN. We discovered a strong positive association between anti-RBD and anti-ECD plasma IgG titer and *in vitro* VN titer.

The data provide important evidence that anti-ECD and anti-RBD IgG titers are a suitable proxy for VN titer. Given the limited availability of VN assays, which are technically complex, require days to set up, run, and interpret, need a biosafety level 3 laboratory when performed with live native SARS-CoV-2 virus, and the relative ease with which ELISA assays can be implemented and performed in a high throughput fashion, we believe our data provide a guidepost for proxy assessments of VN titers relevant to the COVID-19 pandemic.

We found that although both anti-ECD and anti-RBD IgG titers correlate well and significantly with *in vitro* VN, anti-RBD IgG titer had a tendency for a stronger correlation than anti-ECD IgG titer. This finding is consistent with a study showing clustering of VN epitopes in the SARS-CoV-1 RBD domain (20). Neutralizing monoclonal antibodies mapped to a region of RBD that has a critical role in attachment to the host ACE2 receptor. Given that the RBD is also the important region for ACE2 receptor binding for SARS-CoV-2 (21,22), it is not surprising that anti-RBD IgG titers correlate well with VN titers. Importantly, our results show that anti-RBD or anti-ECD IgG titers of 1:1350 discriminated the presence of an adequate VN titer as recommended by the FDA for COVID-19 convalescent plasma with a probability of 80%. Future studies are required to determine if a VN titer of ≥1:160 has therapeutic benefit. Regardless, our findings clearly indicate that an anti-RBD IgG titer cutoff can be established that serves as a suitable proxy for VN titer.

Our findings greatly expand on recent work showing a relationship between anti-S ELISA and microneutralization titer in 9 samples using a 48-hour post-infection microneutralization assay assessing “whole-well” optical density (11). Suthar et. al. have also demonstrated that RBD-specific IgG endpoint titer correlates well with a focus-reduction neutralization assay (10). Li et al. reported a positive correlation between SARS-CoV-2 VN titer and S-RBD–specific IgG titer, with a serum VN titer of 1:80 as approximately equivalent to a titer of 1:1280 for S-RBD–specific IgG (23). Because of differences in the VN assay used, their titers and those we report here are not equivalent. Harvala et. al. also reported VN and anti-spike ELISA titers were correlated (24), although there were several differences between that study and ours. For example, all donors were male, plasma was collected >28 days after symptom resolution, a different virus strain was used, no repeat donors were studied, and the association with clinical symptoms was not assessed. In addition, they did not study samples obtained during community screening of asymptomatic individuals. Herein, we compared results from two independent neutralization assays run blinded in two independent laboratories. The traditional VN assay (assay one) assessed protection from SARS-CoV-2 virus infection as determined by presence of cytopathic effect three days post-infection. In contrast, assay two (VN2) determines the percentage of SARS-CoV-2 virus-infected cells 24 hrs post-infection as a measure of early virus replication and susceptibility to host-cell infection. These two different approaches to VN assessment, and the robustness of the correlation between the results of the two different assays, adds confidence to our conclusion that anti-RBD IgG and anti-ECD IgG titers measured by ELISA serve as a very reliable surrogate of VN.

Of particular note, approximately one-third of convalescent plasma donors in our study did not meet the FDA-recommended cutoff of 1:160 for VN titer. This finding is consistent with the 60% that did not meet the target neutralization threshold of 1:100 recently described in the Harvala study (24). However, the inability to directly compare titers between laboratories highlights an unmet need for the development of international standards to enable comparisons of SARS-CoV-2 serological assays between laboratories. An increasing number of COVID-19 patients are being treated globally with convalescent plasma. For example, under an FDA-approved expanded access protocol, 20,000 transfusions have already occurred in the United States alone (6). Inasmuch as convalescent donor plasma likely will continue to play an important role in treatment of COVID-19 patients as efforts are made to manufacture polyclonal hyperimmune immunoglobulin and neutralizing monoclonal antibodies, we felt it necessary to determine if certain donor characteristics may associate with high VN titer. We found that antibody titers were associated with disease severity and hospitalization status. Among all COVID-19 symptoms and donor characteristics assessed, the presence of dyspnea was the best symptom to discriminate the presence of an adequate IgG antibody titer. Although the sample size is small, we found that even for donors who donated plasma twice/week for up to seven donations, there was no significant decrease in titers as assessed by the IgG ELISAs and VN. We believe these data could inform efforts to recruit plasma donors for therapeutic purposes. The finding that increased COVID-19 disease severity is associated with a more robust humoral immune response is consistent with previous studies of SARS and dengue hemorrhagic fever patients (25), but contrasts with a recent report analyzing COVID-19 patients (26). It is possible that differences in antibody testing platforms could account for the contrasting observations. The list of emergency use authorized antibody testing platforms is rapidly expanding, and test performance, especially as it relates to virus neutralization, will be important to understand (27).

Analysis of the available genomes for the SARS-CoV-2 strain pairs infecting convalescent donors and recipients found few differences in the inferred amino acid sequences, and no association between magnitude of humoral immunity, disease severity, or infecting strain genotype. Because our sample size is small, more work is required in this area.

Several important matters remain unanswered with respect to anti-S protein IgG antibodies. First, although many believe, and some experimental animal infection data support (28), that antibodies directed against S protein confer protection from SARS-CoV-2 infection or reinfection, this remains unproven in humans. Second, although our data and work by others show a strong relationship between anti-S protein IgG titers and *in vitro* VN, it will be important to determine if IgG antibody titer against this protein is a significant correlate of protective immunity in humans. This is an especially important topic given the massive efforts globally on using S protein as a vaccine.

### Limitations

Our study has several limitations. The study was retrospective, only IgG titers were analyzed, and all VN studies were conducted *in vitro*. Plasma from the convalescent donors was used for VN assays, whereas serum samples were used for ELISA assays. As such, the findings may not be entirely applicable to all antibody testing platforms or other sample types.

### Conclusions

Taken together, the data clearly show that anti-RBD and anti-ECD IgG titers serve as important surrogates for *in vitro* VN activity. A substantial fraction of convalescent plasma donors may have VN titers below the FDA recommended cutoff of ≥1:160. Dyspnea, hospitalization, and higher disease severity were associated with higher VN titer. Importantly, a small percentage of asymptomatic individuals have virus-neutralizing antibodies, including some with a titer of ≥1:160. In the aggregate, it is reasonable to think that our findings provide impetus for widespread implementation of anti-RBD and anti-ECD IgG antibody titer testing programs. The resulting data could be useful in several settings, including, but not limited to, identification of plasma donors for therapeutic uses (e.g., convalescent plasma transfusion and/ or source plasma for fractionation in the manufacture of hyperimmune globulin) (5, 11), assessment of recipients of candidate vaccines, assessment of recipients of passive immune therapies, assessment of previously infected individuals, and identification of asymptomatic individuals with antibody against SARS-CoV-2.

## Methods

### Convalescent Plasma Donors

Convalescent plasma was obtained by apheresis using the Trima Accel automated blood collection system (Terumo BCT) and processed by standard blood banking protocols under Houston Methodist human subjects protocol PRO00025121. FDA recommendations for COVID-19 convalescent plasma donor collection were followed (5). Each donor had laboratory-confirmed SARS-CoV-2 infection based on a positive RT-PCR test. All plasma was donated by recovered and healthy COVID-19 patients who had been asymptomatic for more than 14 days. Donors were between 18-65 years old. All donors provided written informed consent and tested negative for SARS-CoV-2 at the time of plasmapheresis. If eligible according to standard blood donor criteria, donors were enrolled in a frequent plasmapheresis program. Donors were documented to be negative for anti-HLA antibodies, hepatitis B, C, HIV, HTLV I/II, Chagas disease, WNV, Zika virus, and syphilis per standard blood banking practices. Disease symptoms (fever, chills, productive or non-productive cough, dyspnea, fatigue, myalgias, headache, runny nose, sore throat, nausea, vomiting, diarrhea, abdominal discomfort, loss of smell or taste, and other), disease severity, hospitalization requirement, and hospitalization course were assessed for each donor. A severity score was assigned as follows: 0 = asymptomatic; 1 = mild disease without dyspnea; 2 = moderate disease with dyspnea that did not require hospitalization; 3 = moderate disease with dyspnea that required hospitalization; 4 = severe disease that required supplemental oxygen; 5 = critical disease that required intensive care unit admission and/or intubation/mechanical ventilation. An aliquot of convalescent plasma product was used for virus microneutralization assays.

Studies were conducted with the approval of the Houston Methodist Research Institute ethics review board, and with informed patient or legally-authorized representative consent when applicable.

### Asymptomatic Donors and VN titers

Samples from asymptomatic individuals were obtained from volunteers screened through an IRB-approved community surveillance protocol (manuscript in preparation). Analysis of 2,814 asymptomatic adults found that 67 (2.4%) had an anti-ECD and anti-RBD IgG ELISA titers of ≥1:150. Sixteen of these 67 specimens were chosen for VN analysis based on the range of titers, including 1:150 (*n* = 5), 1:450 (*n* = 5), and 1:1350 (*n* = 6) (Table S1).

### Specimens from SARS-CoV-2 Naïve Donors

Ten naïve human plasma specimens (negative controls) were obtained from samples biobanked in Houston well before SARS-CoV-2 was described in China, the United States, or elsewhere.

### RT-PCR Testing for SARS-CoV-2 Infection

Symptomatic patients with a high degree of suspicion for COVID-19 disease were tested in the Molecular Diagnostics Laboratory at Houston Methodist Hospital using an assay filed for under Emergency Use Authorization (EUA) from the U.S. Food and Drug Administration (27). The assay follows the protocol published by the World Health Organization (29) and uses a 7500 Fast Dx instrument (Applied Biosystems) and 7500 SDS software (Applied Biosystems). Testing was performed on nasopharyngeal or oropharyngeal swabs immersed in universal transport media (UTM), bronchoalveolar lavage fluid, or sputum treated with dithiothreitol (DTT).

### SARS-CoV-2 ELISAs

Detailed ELISA methods have been recently described (30). The ELISA used to measure antispike IgG antibodies in donor serum specimens was performed as follows. Briefly, ECD purified recombinant protein used comprises amino acid residues 1 – 1208, and the RBD comprises amino acids 319 - 591 of SARS-CoV-2 spike protein (GenBank MN908947). Microtiter plates were coated with either purified recombinant SARS-CoV-2 ECD or RBD. Human mAb CR3022 that targets the receptor-binding domain (RBD) of SARS-CoV (31) was used as a positive control. Negative serum control was included on each microtiter plate. Serial dilutions of serum were added, incubated for 1 h, washed, incubated with goat anti-human IgG Fab HRP (Sigma A0293), and washed. ELISA substrate (1-step Ultra TMB, Thermo Scientific cat# 34028) was added, the plates were developed until the top dilution reached the saturation point, and the reaction was stopped with H2SO4. Plates were read at an absorbance of 450 nm.

A similar ELISA was used to study anti-spike ECD antibody titers in serum obtained from surveilled asymptomatic individuals. Recombinant proteins were produced as described above. All samples were tested with an initial screen assay and IgG antibody titers were subsequently performed on positive samples. For the screening assay, patient serum samples and negative control samples were diluted 1:50 in PBS containing 2% nonfat milk prior to addition to the plate. Patient sera that were identified as positive by the screening assay were subsequently titered by 1:3 serial dilutions in PBS-M to create 1:50, 1:150, 1:450, 1: 1350, and 1:4050 final dilutions. Titer was defined as the last dilution showing an optical density greater than average negative control plus three standard deviations.

### SARS-CoV-2 Microneutralization Assay (VN)

The ability of plasma samples to neutralize SARS-CoV-2 host-cell infection was determined with a traditional VN assay using SARS-CoV-2 strain USA-WA1/2020 (NR-52281-BEI resources), as previously described for SARS-CoV (15). The assay was performed in triplicate, and a series of eight two-fold serial dilutions of the plasma or serum were assessed. Briefly, 100 tissue culture infective dose 50 (TCID50) units of SARS-CoV-2 was added to two-fold dilutions of heat inactivated serum or plasma, and incubated for 1 h at 37°C. The virus and plasma mixture was added to Vero E6 cells grown in a 96-well microtiter plate, incubated for 3 d, after which the host cells were treated for 1 h with crystal violet-formaldehyde stain (0.013% crystal violet, 2.5% ethanol, and 10% formaldehyde in 0.01 M PBS). The endpoint of the microneutralization assay was designated as the highest plasma dilution at which all three, or two of three, wells are not protected from virus infection, as assessed by visual examination.

### SARS-CoV-2 Microneutralization Assay Two

A second SARS-CoV-2 microneutralization assay (VN2) was adapted from an assay used to study Ebola virus (32). This assay also used SARS-CoV-2 strain WA1. Plasma specimens were diluted in cell culture media in duplicate. Serum from naïve and SARS-CoV-2 convalescent individuals was used as negative and positive controls, respectively. Diluted plasma was mixed with the SARS-CoV-2 WA1 strain, incubated at 37° C for 1 h, then added to Vero-E6 cells at a target MOI of 0.4. Unbound virus was removed after 1 h incubation at 37° C and culture media was added. Cells were fixed 24 h post-infection, and the number of infected cells was determined using SARS-CoV-S specific mAb (Sino Biological 401430-R001) and fluorescently labeled secondary antibody. The percent of infected cells was determined with an Operetta high content imaging system (PerkinElmer) and Harmonia software (33). Percent neutralization for each plasma sample at each dilution was determined relative to untreated, virus only control wells.

### SARS-CoV-2 Genome Sequencing and Analysis, and Clade Assignment

Libraries for whole virus genome sequencing were prepared according to version 1 or 3 of the ARTIC nCoV-2019 sequencing protocol (34). Long reads were generated with the LSK-109 sequencing kit, 24 native barcodes (NBD104 and NBD114 kits), and a GridION instrument (Oxford Nanopore). Short reads were generated with the NexteraXT kit and a MiSeq or NextSeq 550 instrument (Illumina). Whole genome alignments of consensus virus genome sequence generated from the ARTIC nCoV-2019 bioinformatics pipeline were trimmed to the start of orf1 ab and the end of orf10 and used to generate a phylogenetic tree using RAxML (https://cme.h-its.org/exelixis/web/software/raxml/indexhtml). Trees were visualized and annotated with CLC Genomics Workbench v20 (Qiagen). SARS-CoV-2 clade assignment was based on procedures described elsewhere (35).

### Statistical Analysis

To assess the correlation between VNs, anti-RBD and anti-ECD ELISA titer data, pair-wise Pearson correlations were performed using the entire dataset, i.e. individuals with single and repeated measurements. To identify the prevalence of donors with VN titers ≥1:160, the frequency distribution of these cases by titer classes critical for RBD, ECD, and VN2 was quantified. Generalized Liner model (GLM), using the first plasma donation data only, was performed between the same variables, as a response, and each of the following predictor factors: dyspnea (yes, no), disease severity (five classes as described above), hospitalization (yes, no) gender (male, female), and age combined into five age groups (<=30, 31-40, 41-50, 51-60 and >60). For variables with more than two factors, a post-hoc t-test (with Bonferroni correction) was used to identify significant pair-wise differences. A linear mixed effect (LME) model was used to analyze the relationship between VNs, anti-RBD, and anti-ECD protein titers, as responses, and days since symptoms, as numerical predictors. Here, we used the whole data set and included the individual’s ID as the random factor, to consider multiple sampling. A similar analysis was used for duration of symptoms but using GLM and selecting only the cases at the first visit. Analysis were performed using log2-trasformed numeric data and the R statistical computing platform (http://www.R-project.org, v. 4.0.0).

## Author Contributions

Project concept (ES, VK, JMM); acquired data (ES, SVK, PAC, TNE, XY, PZ, ZJ, SWL, RJO, JC, BC, DMT, JL, JDG, JC, GCI, RHN, IMB, DG, RMR, SS, IBP, IMC, NJ, LP, KH, AH, JD, VK); analyzed data (VK, SVK, IMC, JMM, ES, PAC); wrote manuscript (ES, VK, JMM); prepared figures (ES, SVK, IMC, VK); provided scholarly advice (CL, PJH, DWB). All authors revised the manuscript and gave final approval for publication.

## Acknowledgments

We are deeply indebted to all of our volunteer plasma donors for their time, their generous gift, and their solidarity. We thank Katharine G. Dlouhy, Curt Hampton, and their team of coordinators and recruiters for outstanding efforts; and Monisha Dey, Cheryl Chavez-East, John Rogers, Ahmed Shehabeldin, David Joseph, Guy Williams, Karen Thomas, and Curt Hampton who were instrumental in efficiently managing the donor center; Drs. Jessica Thomas and Zejuan Li, Erika Walker, the very talented and dedicated molecular technologists, and the many labor pool volunteers in the Molecular Diagnostics Laboratory for their dedication to patient care; the many donor center and blood bank phlebotomists and technologists for their dedication to donor and blood safety; Sasha Pejerrey and Adrienne Winston for editorial assistance; Brandi Robinson, Harrold Cano, and Cory Romero for technical assistance; Claude Moussa, Heather Patton, and the many members of the laboratory information technology team for rapidly implementing the necessary electronic workflows; Pamela McShane, Dilzi Mody, and the many members of the biorepository team for their meticulous management of patient samples; and Christina Talley, Dr. Susan Miller and Mary Clancy for consistent, thorough, and outstanding advice. We express our gratitude to Manuel Hinojosa and Mark Vassallo for their extensive efforts to rapidly procure resources, and Dr. Roberta Schwartz for her efforts in implementing screening of asymptomatic individuals. We are indebted to Drs. Marc Boom and Dirk Sostman for their support, and to many very generous Houston citizens and businesses for their tremendous philanthropic support of this ongoing project, including but not limited to anonymous, Ann and John Bookout III, Carolyn and John Bookout, Ting Tsung and Wei Fong Chao Foundation, Ann and Leslie Doggett, Freeport LNG, the Hearst Foundations, Jerold B. Katz Foundation, C. James and Carole Walter Looke, Diane and David Modesett, the Sherman Foundation, Paula and Joseph C. “Rusty” Walter III, and Aramco Americas. Dr. Jason S. McLellan (University of Texas at Austin) graciously provided the mAb CR3022 and the spike protein expression vectors, and we thank the members of the Center for Systems and Synthetic Biology at the University of Texas at Austin for technical assistance. We thank Terumo BCT for continuously and rapidly supplying blood collection devices and supplies, and Victoria Cavener, Meera Divek and Team COVID-19 serology at Penn State for their timely and generous technical assistance and logistical support.

This study was supported by the National Institutes of Health grants AI146771-01 and AI139369-01, and the Fondren Foundation, Houston Methodist Hospital and Research Institute (to JMM). This research has been funded in whole or part with federal funds under a contract from the National Institute of Allergy and Infectious Diseases, National Institutes of Health, Contract Number 75N93019C00050 (to JL and GCI). A portion of this work was funded through Cooperative Agreement W911NF-12-1-0390 by the Army Research Office (to JDG). We gratefully acknowledge seed funding from the Huck Institutes of the Life Sciences for the studies at Penn State, together with the Huck Distinguished Chair in Global Health award (to VK).

---

**Supplemental Fig. 1.**
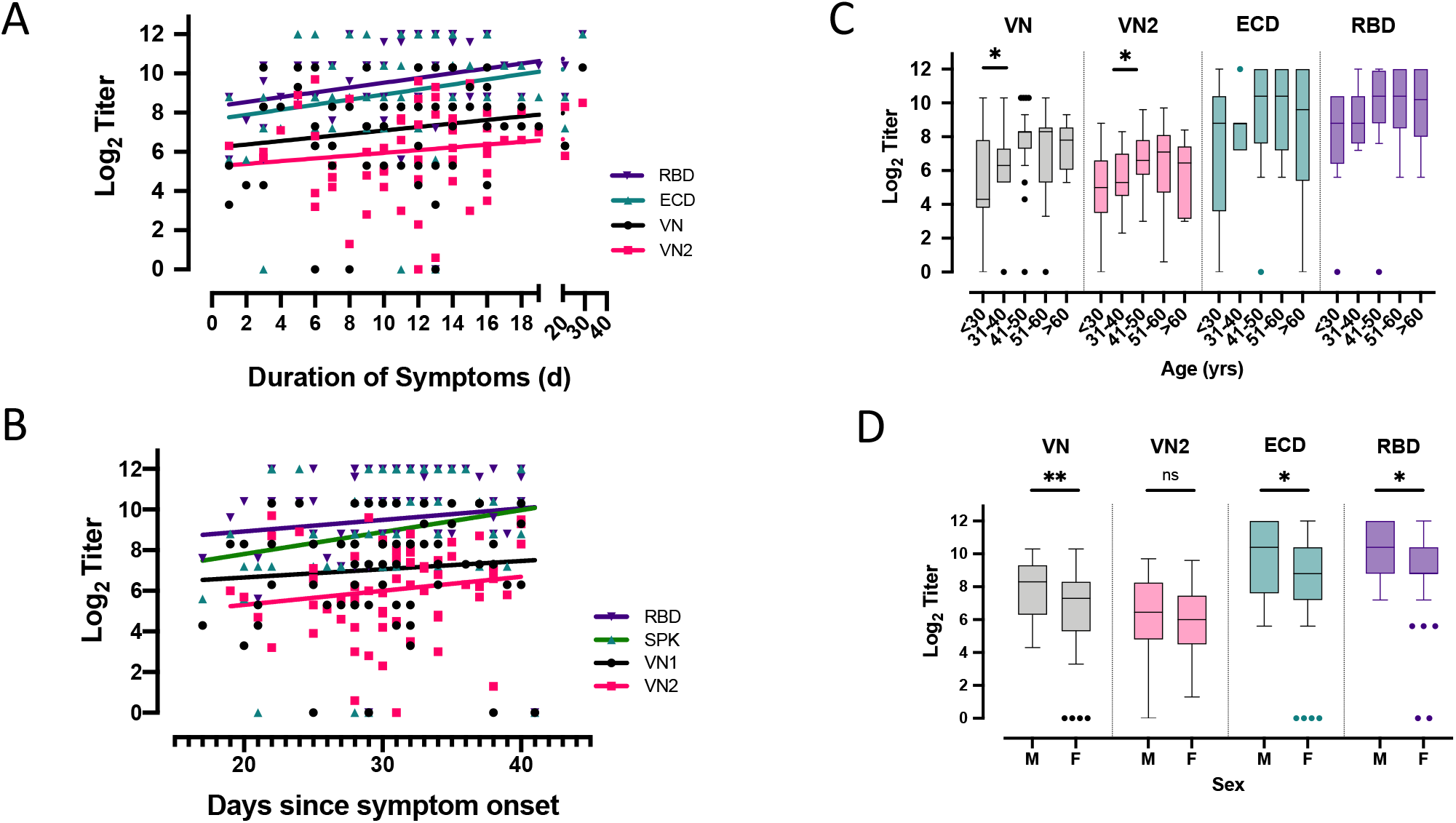
Relationships between VN titer, VN2 titer, anti-ECD IgG titer, and anti-RBD titer by A) Duration of symptoms and B) Days since symptom onset. A linear regression (line) is fitted to each set of data. C) Boxplots of VN, ECD, and RBD by donor age, or D) sex The median, minimum, maximum, first and third quartile and extreme values including pair-wise significant comparisons (*=p<0.05, **= p<0.01) are reported.

**Supplemental Fig. 2.**
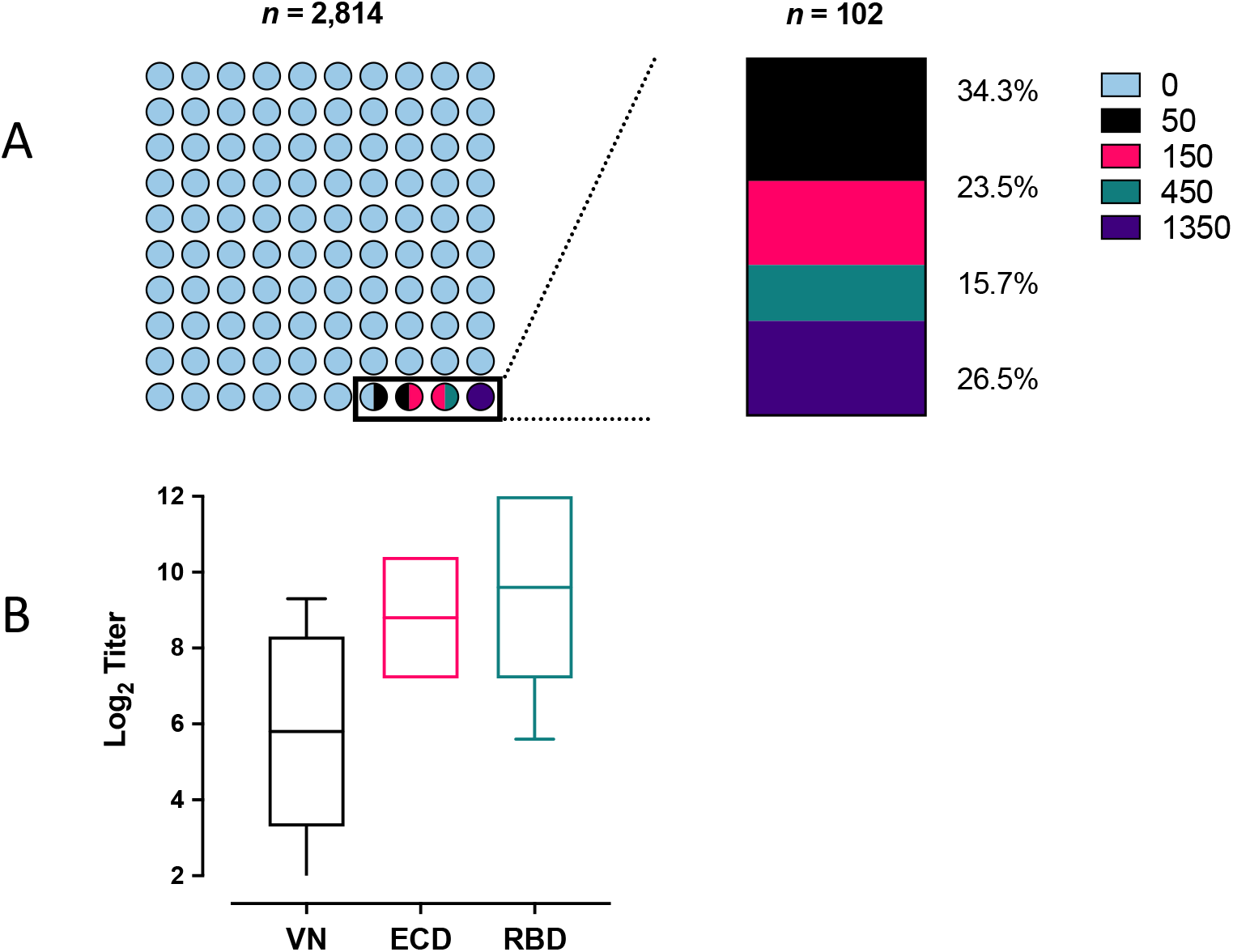
Asymptomatic surveillance sample. A. Counts of ECD titer classes in surveillance screen, left. Percent of individuals in each class are shown, right. B. Boxplots of VN, anti-ECD and anti-RBD titers in a subset of samples (*n* = 16).

**Supplemental Table 1:** Demographics and Characteristics of Asymptomatic Surveillance Subjects Selected for Study.

| Sample # | Subject ID | Gender | Age | ECD (Titer) | RBD (Titer) | VN (Titer) |
| --- | --- | --- | --- | --- | --- | --- |
| 94 | HMH_Surv_01 | M | 20 | 1350 | 1350 | 160 |
| 95 | HMH_Surv_02 | F | 36 | 1350 | 4050 | 640 |
| 96 | HMH_Surv_03 | M | 45 | 1350 | 4050 | 320 |
| 97 | HMH_Surv_04 | F | 42 | 1350 | 4050 | 640 |
| 98 | HMH_Surv_05 | F | 33 | 1350 | 4050 | 320 |
| 99 | HMH_Surv_06 | F | 28 | 1350 | 4050 | 320 |
| 100 | HMH_Surv_07 | M | 49 | 450 | 1350 | 320 |
| 101 | HMH_Surv_08 | M | 38 | 450 | 150 | 10 |
| 102 | HMH_Surv_09 | F | 45 | 450 | 150 | 10 |
| 103 | HMH_Surv_10 | M | 39 | 450 | 450 | 1 |
| 104 | HMH_Surv_11 | F | 25 | 450 | 450 | 40 |
| 105 | HMH_Surv_12 | F | 30 | 150 | 150 | 20 |
| 106 | HMH_Surv_13 | M | 32 | 150 | 1350 | 80 |
| 107 | HMH_Surv_14 | F | 44 | 150 | 150 | 10 |
| 108 | HMH_Surv_15 | M | 31 | 150 | 50 | 1 |
| 109 | HMH_Surv_16 | F | 33 | 150 | 450 | 1 |

Abbreviations: F, Female; M, Male; ECD, anti-spike ectodomain; RBD, Receptor Binding Domain; VN, Virus Neutralization

